# An improved method for the expression screening of membrane protein-GFP fusions in yeast

**DOI:** 10.1101/172114

**Authors:** Darren Baldock, Judith Sheldon, Ravi Tailor, Katherine Green, John Ray, Shradha Singh, Kathryn Brocklehurst

## Abstract

The expression and purification of membrane proteins is an extremely challenging area of work within Protein Science. Membrane proteins are required for compound screening and structure determination in industry. Here we describe some new and innovative methodology in developing the membrane protein GFP fusion primary expression screening in yeast. This methodology enables the expression of membrane proteins fused to GFP in both *Saccharomyces cerevisiae* and *Pichia pastoris* systems. This capability helps facilitate screening of constructs to establish which are suitable for membrane protein production for compound screening and structure determination

In terms of the primary screening work, we have developed both agar plate and liquid plate expression methodology in yeast. The two approaches correspond well, but the agar plate method is more rapid and we have shown it to have the advantage of allowing cells to be taken directly into confocal microscopy for immediate cell localisation data. Innovative work to extend the methanol induction time in the *Pichia* agar plate method established good differentiation from the background. A novel agar plate method was also developed for *S.cerevisiae* which is also presented. These screening methods allow triaging of constructs for either membrane protein preps for biochemical assays or progression to fluorescence size exclusion chromatography; where various detergents can be screened to determine the most appropriate for membrane protein solubilisation, the starting point for purification, crystallisation and structure determination.

Membrane targets depicted to demonstrate the improved primary screening methodology are a copper transporter Ctr1p from *S.cerevisiae* and a water transporter Aqp4 from human origin.

**Highlights:** An improved method for the production of recombinant MP-GFP fusions in yeast is presented using agar plates.

An agar plate method for MP-GFP expression screening is described for *Pichia pastoris*, with improved induction methodology by the simple addition of methanol, allowing longer induction times for expression clarity.

A new simple rapid agar plate method for MP-GFP expression screening is described for *Saccharomyces cerevisiae*.

Cells can be taken directly from agar plates into confocal microscopy studies for immediate cell localisation data and triaging.

Liquid plate based screening methods are also described for both yeasts in comparison, to show there is corresponding data, helping validate the new agar plate methods.

## 1. Introduction

It has been estimated that membrane proteins account for approximately 20-30% of the genes coded in all sequenced genomes [1], including the human genome [2]. Given their critical role in many biological processes, for example transport of metabolites, energy generation and as pesticide and drug targets, there is intense interest in determining their three dimensional structure. However compared to soluble proteins, membrane proteins are highly under-represented in the Protein Data Bank. In fact, of the >100,000 released structures in the PDB (<u>http://www.rcsb.org/pdb/</u>) less than 1000 (<u>http://blanco.biomol.uci.edu/mpstruc/</u>) are classified as membrane proteins. This reflects the technical difficulties of working with membrane proteins which are naturally expressed at relatively low levels; rhodopsin is one of a few exceptions [3, 4]. The production of recombinant membrane proteins for structural and functional studies remains technically challenging, also due to low levels of expression. Over-expression of recombinant membrane proteins in heterologous cells often results in misfolding and poor targeting to the membrane which limits the overall level of production. Downstream, the inherent instability of many membrane proteins once solubilized in detergents is also problematic.

The yeast expression systems were the first successful system for recombinant expression of eukaryotic integral membrane proteins (IMPs) for crystallographic studies. [5, 6, 7] The most common yeast strains for the over-expression of IMPs for structural studies are the baker’s yeast, *Saccharomyces cerevisiae* (*S.cerevisiae*) [5, 6] and *Pichia pastoris* (*P.pastoris*) [8, 9, 10, 11], the methylotrophic yeast, able to utilise methanol as the sole carbon source. Both strains are cost effective eukaryotic expression systems used in many academic and industrial laboratories globally. Expression in *S.cerevisiae* uses a multicopy plasmid system while in *P.pastoris* an integrated vector system is used.

Most vectors used in *S.cerevisiae* expression systems are propagated using the high copy 2-micron plasmid replication origin [12]. Selectable markers are used in *S.cerevisiae* transformation. The most commonly occurring selective markers are the Ura3, Leu2, Trp1 and His3 [13]. Most engineered *S.cerevisiae* expression strains are designed with one or more of these 3 genes knocked out. In this study, the commercial yeast strain, BY4741 is used and originally acquired from the ATCC, the yeast strain repository. It is a yeast knockout strain with the following phenotype: MATa, his3∆1, leu2∆0, met15∆0, ura3∆0. This allows for its use as an expression host using his, leu or ura as auxotrophic markers. In this study we use this strain in conjunction with the pYES2 plasmid vector from Invitrogen carrying your gene of interest (GOI). This vector has a URA3 gene. As the host strain, BY4741 is unable to manufacture its own uracil, therefore only colonies are produced which have successfully taken up the vector will grow on minimal SD (synthetic defined) media which contains all key nutrient supplements minus uracil. This is a very good method of selection with no possibility of false positives. The vector also contains the GAL1 promotor which allows for galactose induced expression. Recombinant expression in *S.cerevisiae* is controlled by the carbon source. During growth and selection, glucose is used as the carbon source. It is only when the carbon source is exchanged for galactose that expression will be induced. This is very tightly regulated with no leaky expression.

There are comparatively a more limited number of vectors for *P.pastoris* expression but the most widely used, pPICZ, uses a promotor derived from the alcohol oxidase I (AOXI) gene, which is inducible by methanol and carries a simple antibiotic based selection system (Zeocin). The *P.pastoris* vector carrying your GOI, integrates into the host cell genome to produce a stable expressing clone. It is not possible to control the number of copies which integrate and so the optimal clone must be experimentally determined. Copy number can be estimated through colony screening in the presence of increasing concentration of antibiotic, Zeocin. However copy number does not necessarily correlate with higher membrane expression levels or functional protein, so it is not sufficient to identify clones based on copy number alone. The *P.pastoris* strain, GS115 (his4) is the most successful for the production of membrane proteins for structural studies and is used in this study.

An increasingly common screening approach is to produce membrane proteins as GFP fusions with expression detected by GFP fluorescence in a primary screen [14, 15, 16, 17, 18]. This enables the construction and expression screening of multiple membrane protein/variants to identify candidates suitable for further investment of time and effort. The GFP reporter is also then used in a secondary screen for cell localisation by visualizing GFP fluorescence following confocal microscopy. Membrane proteins that show both a good expression level and membrane localisation, with minimum degradation as indicated by the absence of free GFP in the cytosol, can be selected for further downstream analysis and further screening.

In this article we describe some new and innovative methodology in developing the membrane protein GFP fusion primary expression screen in yeast. Original solutions have appeared in all stages of agar plate culturing methodologies (the overall screening strategy is presented in Fig1).

**Fig1.**
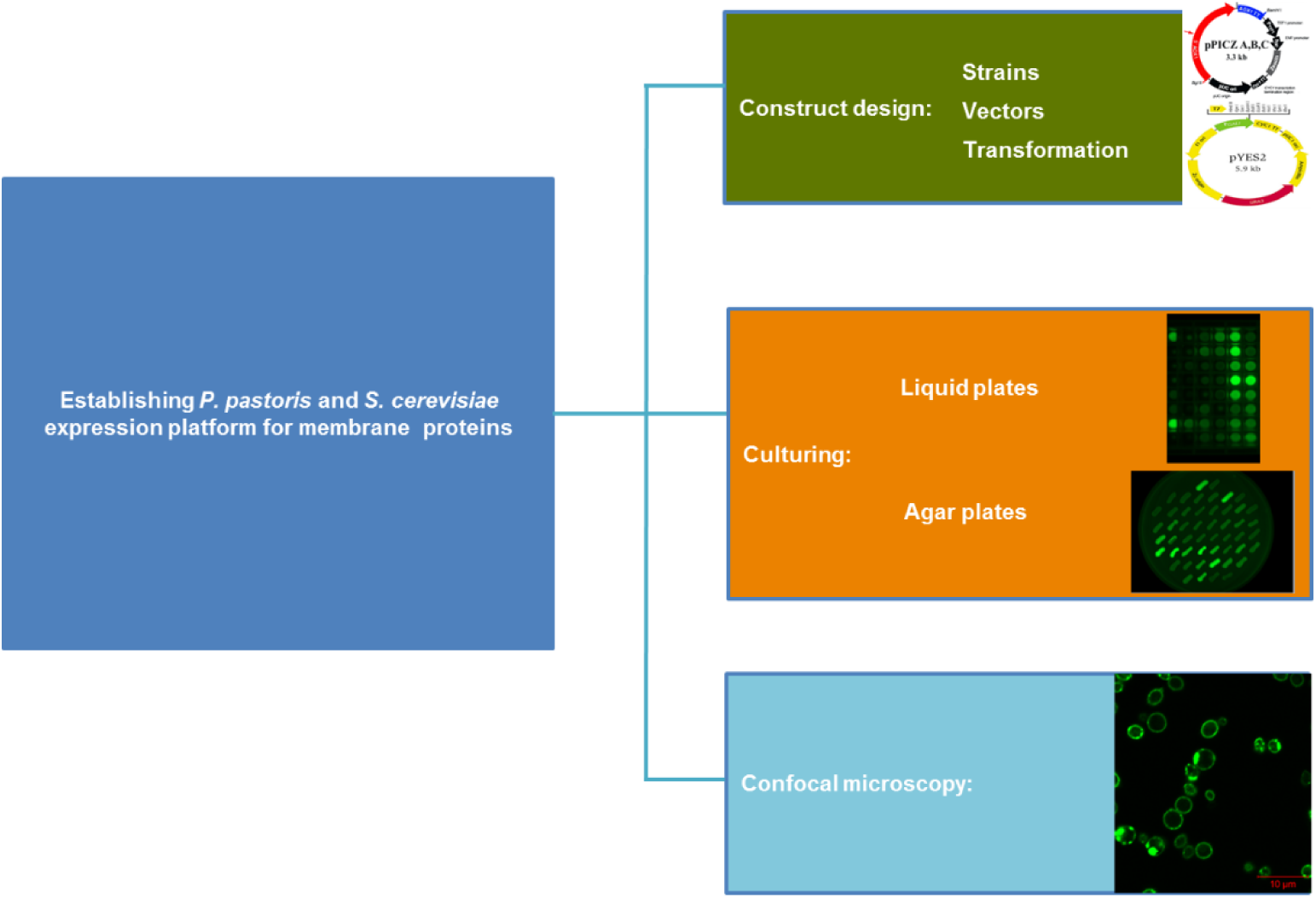
Screening strategy: Schematic diagram to show the overall strategy adopted to establish membrane protein expression screening in yeast.

Membrane targets depicted to demonstrate the improved primary screening methodology are a copper transporter Ctr1p from *S.cerevisiae* [19] and a water transporter Aqp4 from human origin [20].

## 2. Materials and Methods

### 2.1 Cloning and Transformation

#### Strains, vectors and cloning strategy

As expression strains, the strains *P. pastoris* GS115 (his4) and *S. cerevisiae* BY4741 (MATa, his3∆1, leu2∆0, met15∆0, ura3∆0.) were used. GENEWIZ Co. (USA) used for all gene synthesis and cloning inclusive of the Yeast-enhanced green fluorescent protein (yeGFP) (S65G; S72A) sequence. The GenBank accession number for the yeGFP3 sequence is U73901, *S.cerevisiae* Ctr1p 1-126 N terminal deletion sequence (UniProt P49573) and full length hAqp4 sequence (UniProt P55807) were cloned into pPICZ vector and pYES2 vector respectively following codon optimisation for each yeast strain. Plasmids, pPICZ and pYES2, were purchased from Invitrogen Co. (Carlsbad, CA, USA). The construct design is shown in Fig2 for insertion into the plasmid vectors.

**Fig2.**
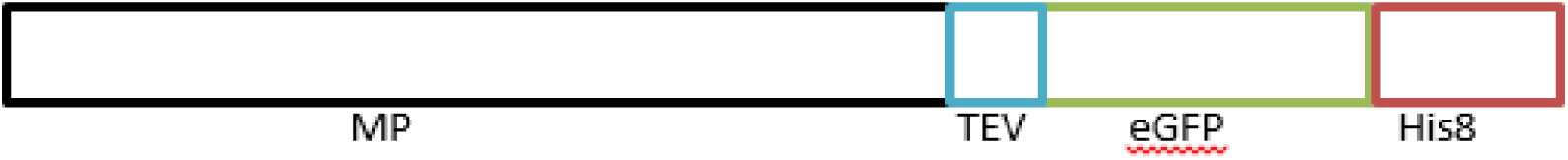
Construct design: The MP constructs (either Ctr1p 1-126 N terminal deletion or full length hAqp4) contain a C-terminal tobacco etch virus protease site (TEV), yeast enhanced green fluorescent protein (eGFP) and a C-terminal eight histidine tag (8His).

#### Making electrocompetent *S.cerevisae* cells

Grow 10ml O/N starter culture of required *S. cerevisiae* strain in YPD medium. After 24hrs, measure the OD600 and inoculate 100ml YPD to ~ 1.0 x 10^5^ cells/ml (OD ~ 0.2) and incubate overnight at 30°C, 250rpm. In the morning measure the OD600, (usually ~4). Perform cell count (see appendix for best method of performing yeast cell counts). Spin down the cells at 1,500 x g. Resuspend the cells in 50 ml: 100mM Li acetate, 10mM DTT (added just before use), 0.6 M Sorbitol, 10mM Tris pH 7.5. Incubate 30mins RT. Centrifuge 3 mins 1500 g 4°C. Discard supernatant. Wash cells by resuspending pellets in 20 ml 1 M ice cold sorbitol. Centrifuge and wash twice more as above. For 5 x 10^7^ cells per transformation resuspend cells to 6.25 x 10^8^ cells/ml in ice cold 1 M sorbitol Freeze cells in 80 µl aliquots. Slow freeze at -80°C in polystyrene. Do not snap freeze. Store at -80°C.

#### Making electrocompetent *P.pastoris* cells [21]

Grow 10ml starter cultures of required *Pichia* strain (GS115) in YPD medium + Ampicillin to prevent bacterial contamination, for 24hrs (~4pm-4pm) at 28°C (increase temp to 30°C in the morning if cultures grew slowly overnight). After 24hrs, measure the OD and use a small amount to inoculate 500ml YPD/Ampicillin to OD = ~0.003 in 3L Thomson flask and incubate overnight at 28⁰C, 250rpm. In the morning measure the OD, which should be ~1 - 1.5. Calculate the number of cells/ml culture using 1AU_600nm_ = 5X10^7^ cells/ml. Calculate what volume you need to harvest: 8X10^8^ cells/transformation, so for 20 transformations you need 1.6X10^10^ cells. Spin down the required volume in multiple falcon tubes for 5mins, 1500g at RT. For example; 4 tubes with cells for 5 transformations in each. Resuspend cells, allowing 8ml per transformation, in: 100mM Li acetate, 10mM DTT (added just before use), 0.6M Sorbitol, 10mM TRIS pH 7.5. Incubate 30mins RT. Centrifuge 3mins 1500g 4°C. Discard supernatants. Resuspend pellets in 1M ice cold sorbitol to 1.5ml per transformation and combine all samples of the same strain for speed, keeping samples on ice between spins. Centrifuge and resuspend twice more as above. Finally resuspend cells to 10^10^ cells/ml in ice cold 1M sorbitol. e.g. for 20 transformations, (1.6X10^10^ cells) resuspend in 1.6ml. Keep cells on ice for use straight away. Freeze remaining cells in 80ul aliquots: slowly at -80°C in polystyrene. Do not snap-freeze.

Linearization of *Pichia* constructs: The vector pPICZA with GOI is linearized with the enzyme Sac I before electroporation.

#### Electroporation

This was performed using the BioRad Gene Pulser Xcell instrument (Cat no 1652660). Place an electroporation cuvette on ice at least 10-15 minutes before performing the transformation. Mix 80ul of competent cells, with 2ug of linearized DNA in the cuvette, mix gently with the pipette and incubate for 5 minutes on ice.

Adjust the electroporation settings as follows: 1500 V, 25uF, 400 Ω. (Cuvette gap = 0.2cm). Place the cuvette in the electroporator chamber and apply the electric pulse. Time constant should read between 9.2-9.6msec. Immediately resuspend the electroporated mixture in 1 ml of ice cold 1 M sorbitol and transfer cells to a sterile tube. Incubate at 30°C, 900rpm for 2hrs then harvest by centrifugation at 2000 g for 10 min. Discard the supernatant and resuspend the pellet in 200ul YPD media.

### 2.2 *Pichia pastoris* agar plate: Growth and Induction plates

Plate out transformants onto YPDS (1% yeast extract [w/v], 2% peptone [w/v], 2% dextrose [w/v], 2% agarose [w/v], 1 M sorbitol) agar plates with 100ug/ml, 500ug/ml and 1000ug/ml Zeocin using 6 well culture plates for 3 days at 30°C. Take single colonies (16 for each Zeocin concentration = 48 for each construct) for induction plates: BMMY with 100ug/ml Zeocin agar plates. Incubate at 30°C.

<u>Novelty</u>: Spike with 100ul methanol in lid 24hrs, 48hrs and 72hrs, incubating plates for a total of 5 days at 30°C. Analyse plates using the BioRad Chemidoc MP imager (Cat no 170-8280) for the presence of GFP (with a blue-light filter to detect the membrane protein-GFP fusion).

### 2.3 *Pichia pastoris* liquid plate: Growth and Induction plates

*P.pastoris* screening in 96 deep well liquid plates with single colonies from YPDS agar plate (1ml cultures). **nb. Use the same colonies as for the agar plate method.** Growth with YPD/Zeocin media (in 0.5ml wells of 96 deep well plate, 900rpm, 30°C, O/N). Use 50ul O/N inoculum to 1ml BMGY/Zeocin deep well plate (make glycerol’s with remaining inoculum). Leave at 30°C, 900rpm for 24 hrs using a VWR Incubating microplate shaker (Cat no 444-7082). Spin down cells and resuspend cells in 1ml BMMY/Zeocin media, leave at 30°C, 900rpm for 24 hrs then the cells harvested by plate centrifugation. Analyse plates using the BioRad Chemidoc MP imager for the presence of GFP (with a blue-light filter to detect the membrane protein-GFP fusion).

### 2.4 *Saccharomyces cerevisiae* agar plate: Growth and Induction plates

Plate out transformants into Growth plates: SD minimal media with –URA DO and 2% glucose agar plates. Incubate at 30°C O/N. Take 8 single colonies for each construct and plate onto Induction plates: SD minimal media with –URA DO and 2% galactose agar plates. Incubate at 30°C for 24hrs.

Analyse plates using the BioRad Chemidoc MP imager for the presence of GFP (with a blue-light filter to detect the membrane protein-GFP fusion).

### 2.5 *Saccharomyces cerevisiae* liquid plate: Growth and Induction plates

*S.cerevisiae* screening in 96 deep well liquid plates with single colonies (1ml cultures). **nb. Use the same colonies as for the agar plate method.** Growth of colony with 1ml SD minimal media with –URA DO and 2% glucose (in wells of 96 deep well plate, 900rpm, 30°C, O/N). Use 50ul O/N inoculum to 500ul SD minimal media with –URA DO and 2% glucose and 100ul 80% glycerol for glycerol’s and freeze -80°C. Agar plate work devised around ref [22], combining membrane protein- GFP fusion fluorescence. Spin down 950ul cells and resuspend cells in 1ml SD minimal media with –URA DO and 2% galactose / raffinose media, leave at 30°C, 900rpm for 24hrs then harvest and analyse plates using the BioRad Chemidoc MP imager for the presence of GFP (with a blue-light filter to detect the membrane protein-GFP fusion).

### 2.6 Confocal microscopy

*P.pastoris* and *S.cerevisiae* cells expressing either eGFP, Ctr1p-eGFP or Aq4p-eGFP are taken directly from either Induction agar plates or induced liquid culture for immediate cell localisation data under confocal microscopy. Twenty microliter of resuspended cells were directly spotted onto microscope glass slides and mounted with a coverslip. Images and GFP fluorescence were acquired using a Carl Zeiss confocal laser scanning microscope (LSM 780) under Argon. GFP was excited at 488nm and emission was recorded at 500-600nm.

## 3. Results

### 3.1 *P.pastoris* expression screening of MP-GFP fusions in agar plate is successfully improved with longer induction times and expression corresponds to liquid plates

The agar plate induction method with methanol shows various clones expressing to varying degrees from GFP fluorescence imaging. It was important to show good differentiation over background. This is important for lowly expressed membrane proteins as the methanol in the agar plate is exhausted after 24hrs. To improve on this methodology, to allow induction to be both monitored for a longer duration (up to 5 days in this study) and to be able to more easily differentiate clear expression over background, a simple addition of 100ul methanol to the lid of the agar plate every 24hrs allowed clarity in data captured. (Fig3). Importantly the data captured from the agar plates is directly comparable to the traditional liquid plate based methods, help validating the data. The agar plate method is more rapid as cells do not need to be grown overnight, require centrifugation or resuspending in induction media, as they do in liquid culture. The cells can be taken from single colonies after transformation plating directly onto Induction agar plates. Using agar plates only requires an incubator, while liquid plate culturing requires liquid handling to dispense media and the requirement for shaking incubators, so agar plate methods are less costly in time, equipment and manual handling procedures.

**Fig3.**
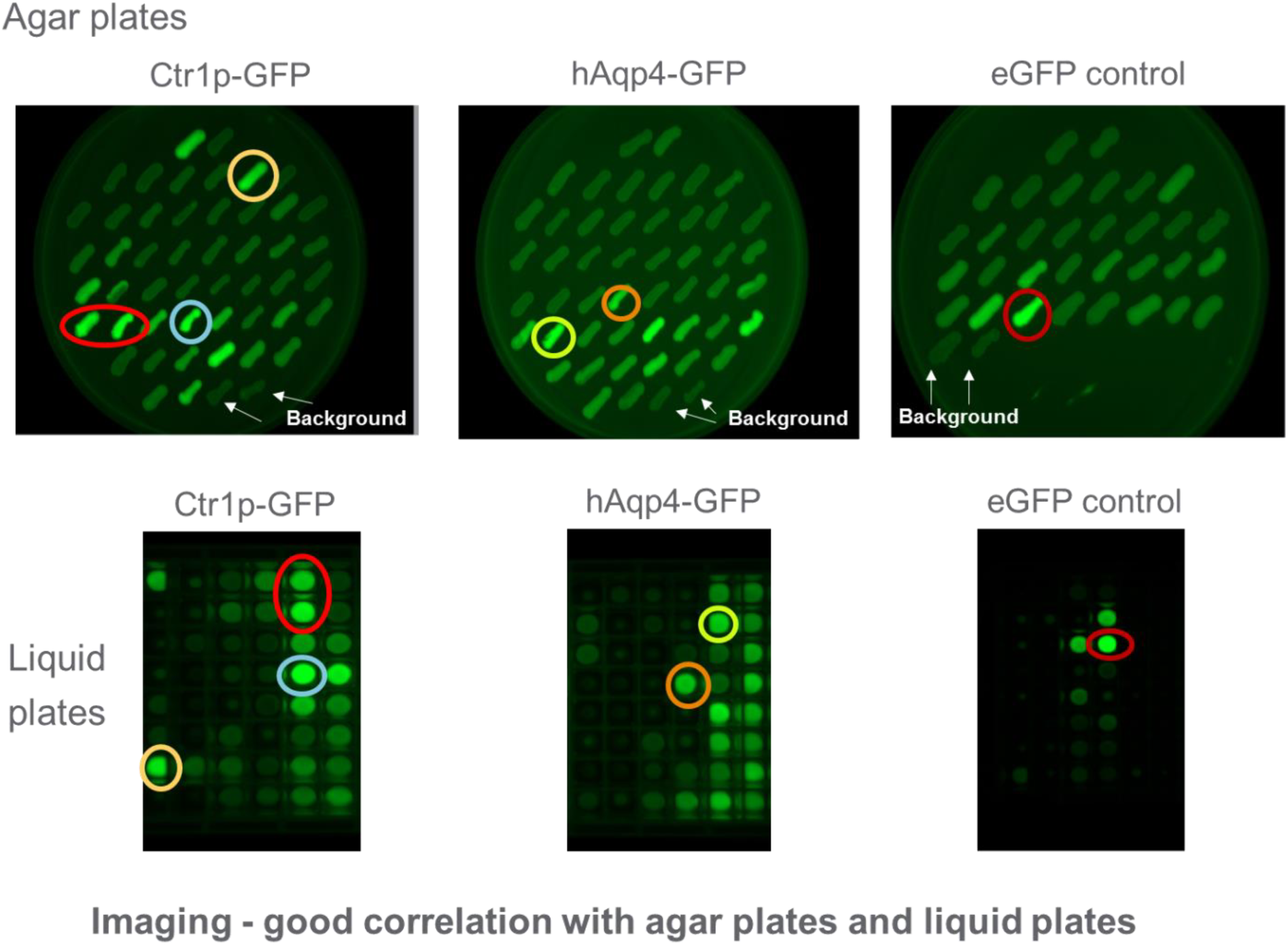
BioRad Chemidoc images detecting GFP fluorescence in *P.pastoris*: Examples of membrane protein–GFP screening (Ctr1p in 1^st^ column, Aqp4 in the 2^nd^ column and eGFP control in the 3^rd^ column) in *Pichia pastoris* using agar plates and liquid plates: Induction agar plates (top row) showing various expression levels of MP-GFP which corresponds to liquid 96 well plate cultures (bottom row).

### 3.2 Agar plate screening allows *P.pastoris* cells to be taken directly and quickly from the plate onto confocal microscopy for immediate cell localisation data

Cells from the agar plate can be immediately taken and examined under confocal microscopy (Fig4). As expected eGFP alone is a soluble protein and is therefore found diffuse throughout the cell cytosol. No GFP fluorescence is observed in untransformed cells. The MP-GFP fusion, Ctr1p, shows a clear ‘green ring’ with patchiness around the plasma membrane, which is typical for Ctr1p. There is also some diffuseness which would imply some free GFP present, which can sometimes indicate a degree of membrane protein degradation [16]. The membrane protein, hAqp4, shows a more specific localisation often observed for Aqp4 and is thought to be ER membrane localised [20].

**Fig4.**
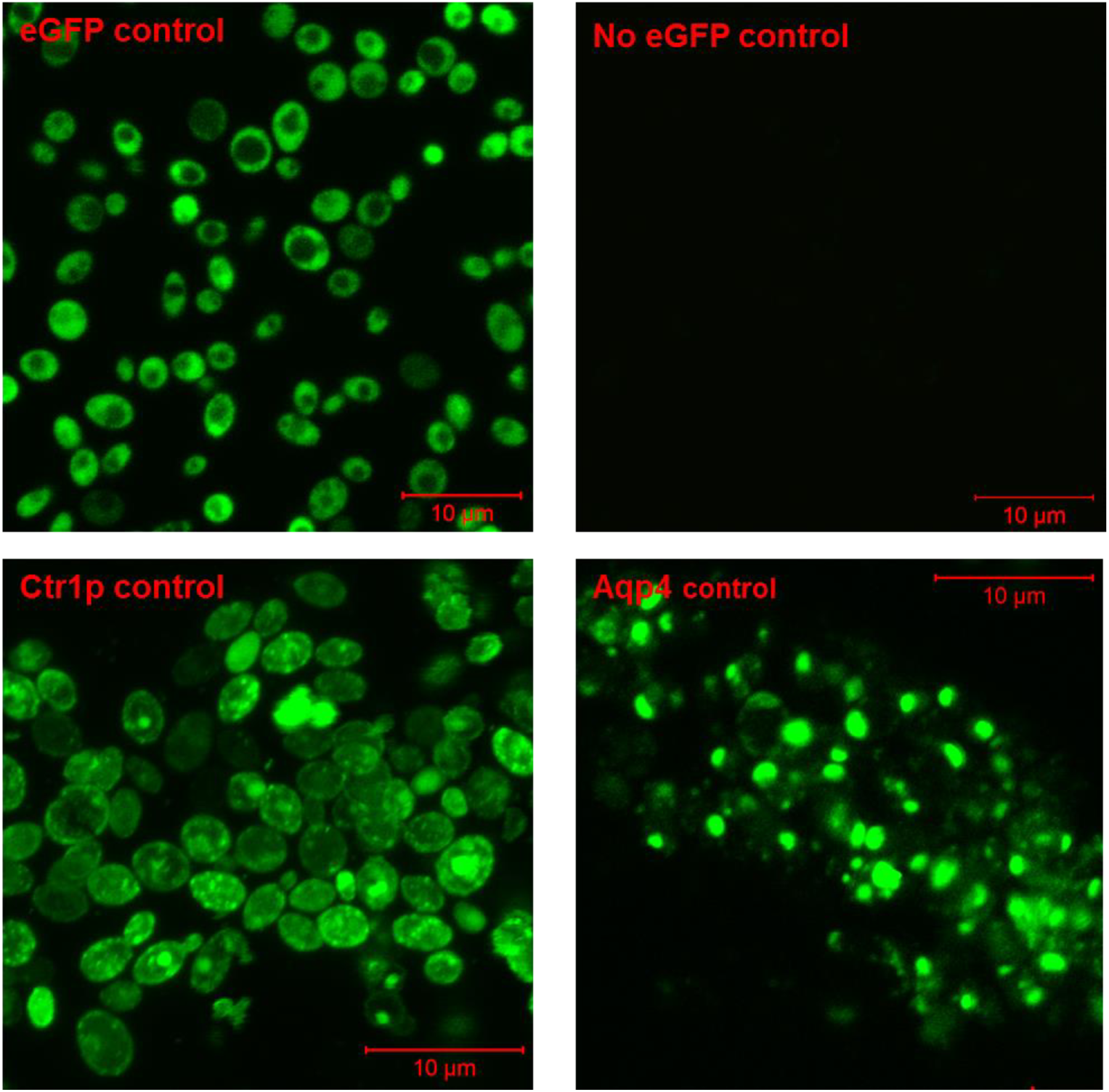
Confocal microscopy and GFP fluorescence: Data of eGFP control (top left panel) and *membrane protein-GFP* control samples (bottom row panels, Ctr1p left, Aqp4 right) from *Pichia pastoris* taken from agar plates. Negative control (top right panel)

An identical confocal pattern is observed with liquid culture samples (Fig5), once again further validating the agar plate based screening method.

**Fig5.**
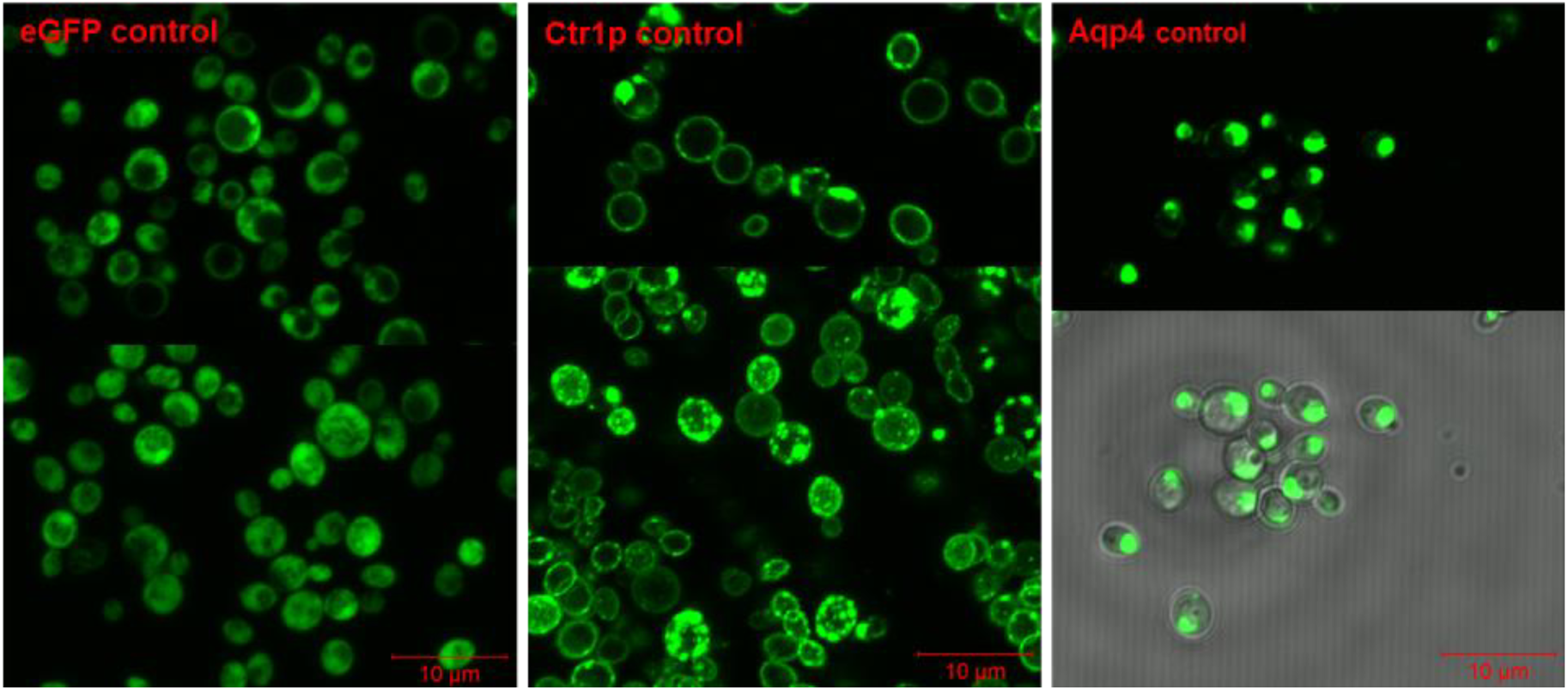
Confocal microscopy and GFP fluorescence: Data of eGFP control (left panel) and *membrane protein-GFP* control samples (middle, Ctr1p and far right panel top, Aqp4) from *Pichia pastoris* taken from liquid plates. Bottom right panel shows merge with white light contrast showing cell location.

### 3.3 Validation of a new *Saccharomyces cerevisiae* agar plate based screening method

The Pichia agar plate method has been previously described [11], where improvements have been made to the induction times in this study. No agar plate method for MP-GFP expression screening in *Saccharomyces cerevisiae* has been described in the literature, apart from liquid culture methods in shake flask [16] and liquid plate [14]. Here we describe a simple new agar plate method which was initially put through a proof of concept study with eGFP (Fig6). Briefly growth/selection plates are made, SD minimal media with –URA DO and 2% glucose agar plates. Induction plates based on using galactose for induction of the Gal1 promotor are made, SD minimal media with –URA DO and 2% galactose agar plates. The concept for this work was based around a publication many years ago describing fluconazole resistance in *Saccharomyces cerevisiae* [22]. As shown in Fig6, a positive proof of concept (POC) study was shown for eGFP in the agar plate method adopted and again this corresponds to liquid plate data.

**Fig6.**
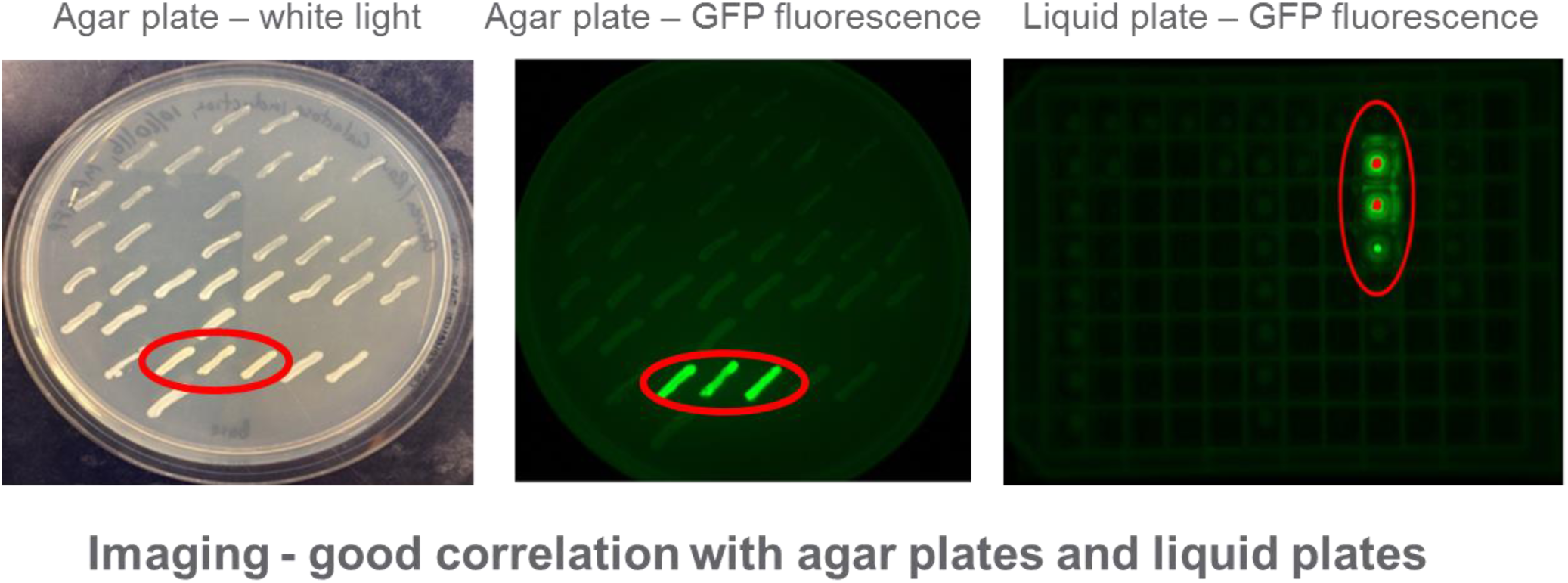
BioRad Chemidoc images detecting GFP fluorescence in *S.cerevisiae*. eGFP screening in *Saccharomyces cerevisiae* using agar plates and liquid plates showing corresponding GFP fluorescence: (left panel shows colonies on plate, GFP fluorescence with middle panel and right panel showing liquid culture plate).

Subsequently this methodology has been tested with the MP-GFP fusion, Ctr1p (Fig7) and shows expression as indicated by GFP fluorescence. No fluorescence is observed in non transformed cells.

**Fig7.**
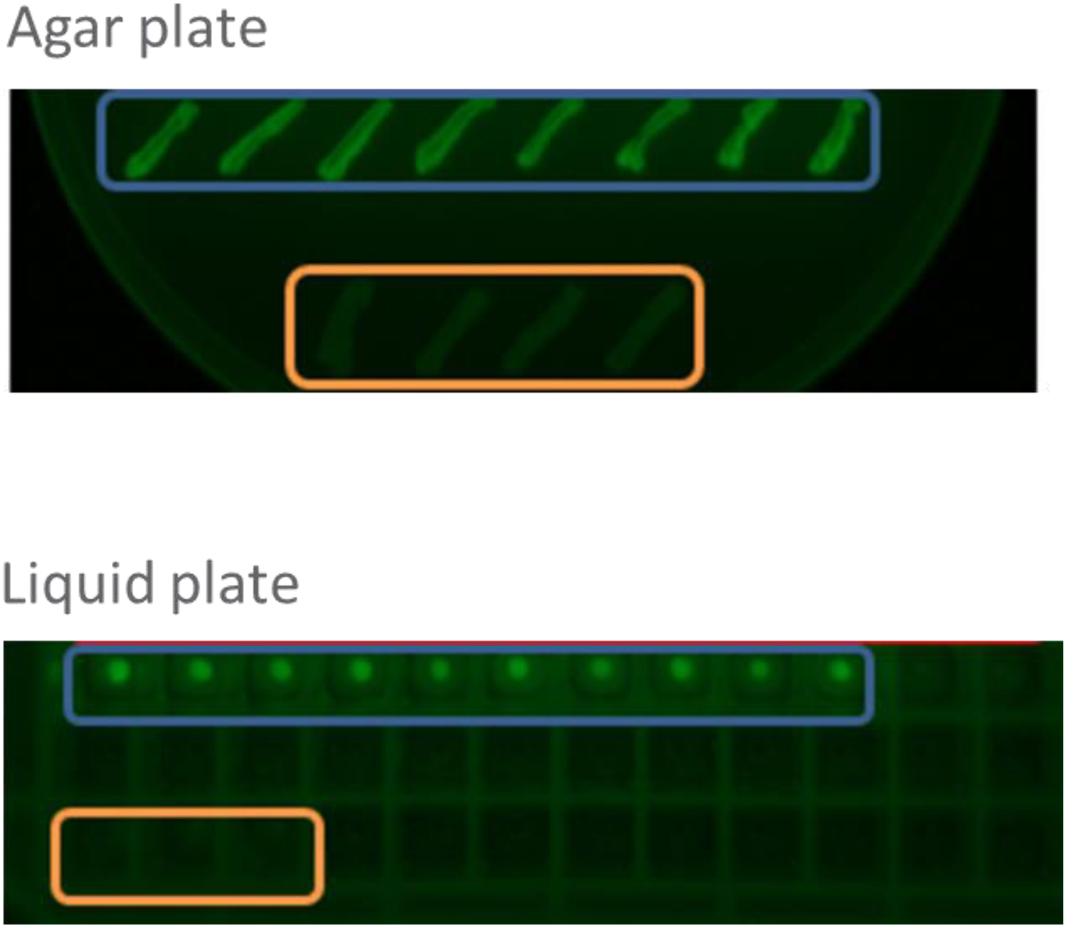
BioRad Chemidoc images detecting GFP fluorescence in *S.cerevisiae*: Membrane protein-GFP screening in *Saccharomyces cerevisiae* using agar plates and liquid plates: Induction agar plate (top panel) showing various expression levels of membrane-GFP fusion, Ctr1p (blue) and no DNA control (brown) which corresponds to liquid 96 well plate cultures (bottom panel).

### 3.4 Agar plate screening allows *S.cerevisiae* cells to be taken directly and quickly from the plate onto confocal microscopy for immediate cell localisation data

As described in section 3.2 for *P.pastoris*, cells can be taken from the galactose induction plates of *S.cerevisiae* expressing eGFP and Ctr1p and examined immediately under confocal microscopy (Fig8).

**Fig8.**
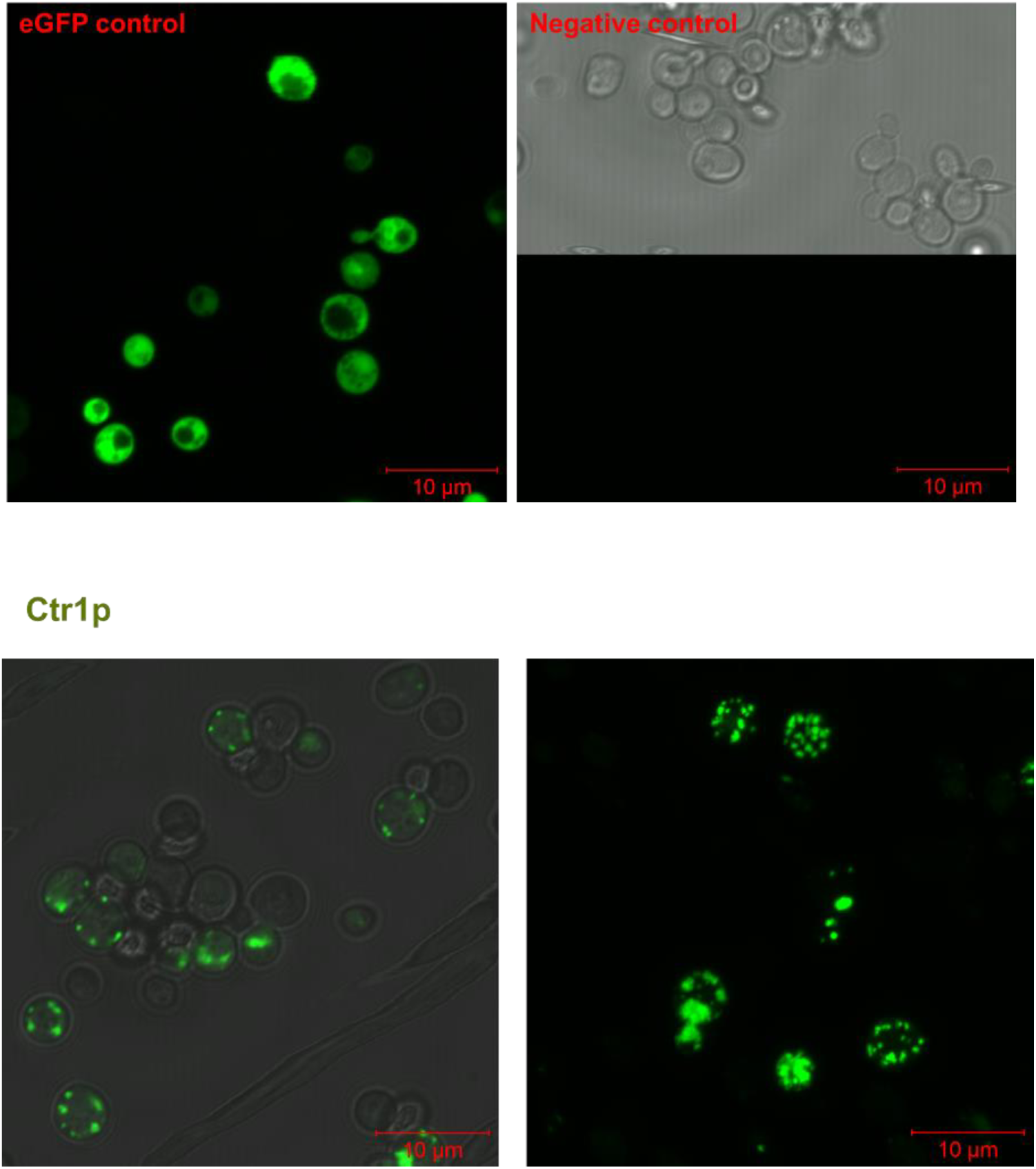
Confocal microscopy and GFP fluorescence: Data of eGFP control (top left panel) and *membrane protein-GFP* control sample, Ctr1p (bottom panel) from *Saccharomyces cerevisiae* taken from agar plates. Negative control (top right panel) shows merge with white light contrast showing cell location.

Confocal data of both eGFP and Ctr1p matches that observed in liquid culture (Fig9), where eGFP is diffuse throughout the cell and Ctr1p is located more around the periphery of the cell albeit more patchy distribution around the plasma membrane [15, 16].

**Fig9.**
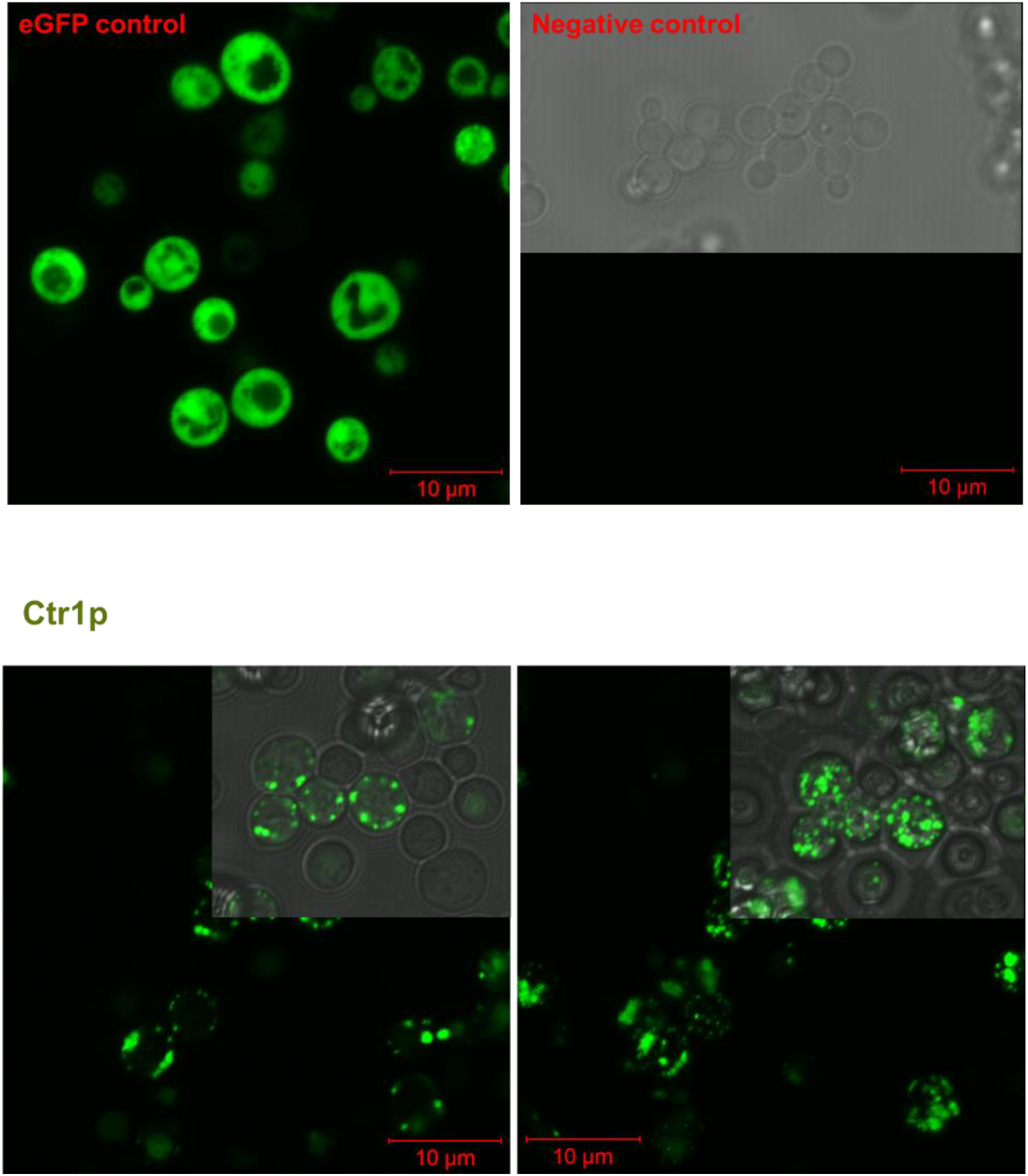
Confocal microscopy and GFP fluorescence: Data of eGFP control (top left panel) and *membrane protein-GFP* control sample, Ctr1p (bottom panel) from *Saccharomyces cerevisiae* taken from liquid plates. Negative control (top right panel) shows merge with white light contrast showing cell location.

## 4. Discussion

In screening for membrane proteins, one requires a method that enables the supply of membrane protein to satisfy the needs for functional assays and structural biology projects. The GFP fusion methodology has over the last 10 years been very successful in achieving this ambition. This article provides further tools to help with MP-GFP screening in yeast.

### 4.1 Agar plate screening and confocal microscopy can help assist MP-GFP expression screening

We have shown successful agar plate screening in both *P.pastoris* and *S.cerevisiae* with membrane proteins Ctr1p, a copper transporter and hAqp4, a water transporter. By taking these cells directly into confocal microscopy this can enhance the primary screen quickly by providing information on which membrane protein constructs are correctly localised to a cellular membrane and the extent of any membrane protein degradation in terms of observing free GFP fluorescence in the cell cytosol. One can then quickly and effectively screen constructs to take further forward for secondary screening which typically involves detergent solubilisation screening and FSEC [18]. *P.pastoris* agar plate screening has been enhanced for MP-GFP screening by the simple addition of methanol to the agar plate lid, enabling extended induction times. This has the great advantage and potential of being able to see some expression of low level expressing membrane proteins and methanol spiking infact mimics what one would do when initially scaling to shake flask expression in *P.pastoris*. Comparing the agar plate and liquid plate screening, it has been shown these are identical in expression (for *P.pastoris* and *S.cerevisiae*), which means the best single colony expressed in liquid culture was the same colony as that used in agar plate screening. The variable expression of different clones, which you expect from *P.pastoris* due to genomic integration differences [11] is also mirrored between the same colonies on the agar and liquid plate methods. With *S.cerevisiae*, being a plasmid based expression there is usually hardly any differences between cells expressing the same membrane protein construct as shown in this study (Fig7). However there are some reports that show you can sometimes get differences which can be attributed to small cell genotypic differences or health status. However like all screening methods the agar plate method will be very valuable for when looking at membrane protein variants e.g. constructs and/or species for example. This method like the liquid screening method testifies itself to medium and large scale screening projects, where stacked plates in a thermal incubator require very little space. The agar plate method is further validated from confocal microscopy data where it is clearly shown that cell localisation data is the same independent of culturing approach, validating the use of the agar screen further. From the confocal microscopy studies, we also observed that the expression of eGFP in controls grown in liquid culture was greater than the expression in agar cultured *Pichia.* (Data not shown). This probably is as a result of the cells being stationary on an agar plate, while in culture there is more aeration and so higher production. Sometimes it is common to see membrane protein ‘patchiness’ around a membrane, however there have been some observations to provide evidence that when solubilised in detergent the protein is more aggregated on FSEC analysis [23], while other studies (R.Owens / L.Bird personal communication) have indicated that a ‘green ring’ around a membrane can also give aggregated protein and this is often attributed due to the detergent solubilisation of the membrane protein into solution and aggregation results from this extraction, rather than the inherent state of the membrane protein in the membrane bilayer. We also noted the interesting observation that not all cells are expressing and some cells are expressing better than others, which no doubt all translates to the ultimate low expression levels observed on membrane protein production in heterologous expression systems.

Infact Histidine and DMSO are commonly used additives to help increase membrane protein expression levels several fold in both *S.cerevisiae* and *P.pastoris* shake flask experimentation [16, 24]. These additives can also be easily added when making up the induction agar plates for yeast, to help further increase expression levels for construct triaging. Industry specific secondary screens in the agar plate method could be developed further with inclusion of target specific inhibitors to ascertain the impact on membrane target expression levels. As some membrane proteins are toxic to cells, there inhibition by compound, may promote the increase in expression level of the membrane protein target. An analogous agar plate induction method is also potentially envisaged with the *E.coli* expression system, for when expressing prokaryotic membrane protein-GFP fusions, whereby an IPTG induction plate method could be adopted.

## 5. Conclusion

There is an overwhelming need in the membrane protein biochemistry and structural biology community to be able to screen for recombinant membrane protein expression in a simple cost effective manner to help triage membrane protein constructs to take further forward for downstream analysis. The first part of the process commonly adopts expression screening by GFP fusion methodology which has greatly enhanced the success of finding good levels of membrane protein expression and protein quality for both functional and structural studies. Here we have described some novel and innovative agar plate methodology that can be easily incorporated into any membrane protein-GFP fusion screening laboratory, working with Yeast. We have shown the methodology to be simple, quick and cost effective, showing comparable data to liquid plate expression screening and with a new approach to further enhance induction for *P.pastoris* agar plate screening. Agar plate screening allows cells to be taken directly off agar plates and placed under confocal microscopy enabling important cell localisation data. A novel *S.cerevisiae* agar plate screening method for MP-GFP fusions has also been described, to help facilitate the ease of membrane protein screening in *S.cerevisiae*, which is a commonly used expression system in many protein science laboratories. Finally, in conclusion, agar plate membrane protein GFP screening data in combination with confocal microscopy data has great potential for triaging membrane protein constructs and variants to help evaluate the best constructs to take further forward for functional assays and structural biology projects.

## Author agreement

There is no conflict of interest of any author in relation to the submission. This manuscript has not been submitted elsewhere for consideration or publication.

## Author contributions

Judith Sheldon performed all the confocal microscopy work; Ravi Tailor helped perform the expression screening experimental work in *P.pastoris* and *S.cerevisae*; Katherine Green helped perform the *P.pastoris* work; John Ray was involved in the molecular biology experimental work; Shradha Singh and Kathryn Brocklehurst helped design the experiments; Darren Baldock planned and designed the experiments, performed the experimental work, analysed the results, wrote the manuscript and prepared all the figures in this manuscript. All authors reviewed the manuscript draft.

## Acknowledgements

We are thankful to Dr Liang Shi for sponsoring this work at Syngenta. We thank Christina Burr for the supply of electrocompetent *S.cerevisae* cells. We are also thankful to Helen Carter for useful discussions around *S.cerevisiae* transformation and Sheila Attenborough for providing support in the writing of this manuscript.

